# NeuriteNet: A Convolutional Neural Network for determining morphological differences in neurite growth

**DOI:** 10.1101/2021.03.31.437885

**Authors:** Joseph Vecchi, Sean Mullan, Josue Lopez, Marlan R. Hansen, Milan Sonka, Amy Lee

## Abstract

**Background:** During development or regeneration, neurons extend processes (*i*.*e*., neurites) via mechanisms that can be readily analyzed in culture. However, defining the impact of a drug or genetic manipulation on such mechanisms can be challenging due to the complex arborization and heterogeneous patterns of neurite growth *in vitro*.

**New Method:** NeuriteNet is a Convolutional Neural Network (CNN) sorting model that uses a novel adaptation of the XRAI saliency map overlay, which is a region-based attribution method. NeuriteNet compares neuronal populations based on differences in neurite growth patterns, sorts them into respective groups, and overlays a saliency map indicating which areas differentiated the image for the sorting procedure.

**Results:** In this study, we demonstrate that NeuriteNet effectively sorts images corresponding to dissociated neurons into control and treatment groups according to known morphological differences. Furthermore, the saliency map overlay highlights the distinguishing features of the neuron when sorting the images into treatment groups. NeuriteNet also identifies novel morphological differences in neurites of neurons cultured from control and genetically modified mouse strains.

**Comparison with Existing Methods:** Unlike other neurite analysis platforms, NeuriteNet does not require manual manipulations, such as segmentation of neurites prior to analysis, and is more accurate than experienced researchers for categorizing neurons according to their pattern of neurite growth.

**Conclusions:** NeuriteNet can be used to effectively screen for morphological differences in a heterogeneous group of neurons and to provide feedback on the key features distinguishing those groups via the saliency map overlay.

**Highlights:**

- NeuriteNet is a machine learning model developed to identify differences in control and experimental groups of cultured neurons based on morphological criteria.
- NeuriteNet’s saliency map highlights the features of the image that associate the neuron with a particular group.
- NeuriteNet outperforms trained researchers in assigning neurons to control or experimental groups with a known morphological difference as well as those with no previously described difference.

## 1. Introduction

In order to process diverse forms of information, neurons extend processes (i.e., neurites) from their soma in patterns that are sometimes quite complex. A variety of assays involving cultured neurons *in vitro* are used to study the mechanisms that regulate neurite growth and branching during development and in response to injury or disease. This type of analysis is relevant for research aimed at treating or preventing peripheral neuropathies (1,2), directing neurite growth toward a desired target (3), and uncovering genes involved in neuropsychiatric/neurodevelopmental diseases (4,5). However, the heterogeneity of neurons isolated from the central and peripheral nervous systems and their complex morphology complicate efforts to identify the impact of experimental manipulations on neurite growth in culture. Large sample sizes and dramatic alterations in morphology are often required to identify a significant effect of an experimental manipulation on neurite growth properties.

While a variety of strategies are available for quantitative analysis of neurite growth, they are not without shortcomings. For example, methods like Scholl analysis (6) require significant manual effort and thus are time-intensive, prone to bias, and results can vary pending investigator experience. These issues can be overcome by automated approaches (7-11), which still require significant pre-processing of images that are to be analyzed and often need to be calibrated with manual procedures. Moreover, the kinds of output generated by automated procedures are generally pre-determined metrics such as the number of branchpoints and terminals or lengths of primary and secondary neurites. Subtle, yet biologically significant, morphological features resulting from an experimental manipulation may be missed with this type of analysis.

To overcome these limitations, machine learning approaches offer promising alternatives for sorting neurons into sub populations based on their morphology (12,13), classifying their dendritic structure (14,15), as well as distinguishing cancer cells from healthy cells (16). Procedurally, these approaches are advantageous as manual manipulations and other user-dependent measurements are not needed. In particular, convolutional neural networks (CNNs) proven successful in image classification tasks for many different image modalities (16). Here, we describe a novel CNN model, NeuriteNet, that classifies images of neurons based on their patterns of neurite growth. The CNN utilizes a branching structure to enable consideration of different spatial resolution levels while learning features of the neurons. This approach allows differentiation of the target categories (*e*.*g*., genotype, treatment, sex, etc.) based on global and local features of the image. We present evidence that NeuriteNet can accurately sort images of neurons into their respective categories based on features extracted from their neurite growth patterns, with a saliency map overlay identifying areas of the images that were used for the sorting.

## 2. Materials and Methods

### 2.1 Animals

All procedures involving animals were conducted in accordance with the NIH Guide for the Care and Use of Laboratory Animals and were approved by the University of Iowa Institutional Animal Care and Use Committee. All mice were maintained on a C57BL/6 (Envigo) background, housed in groups on a standard 12:12 hour light: dark cycle with food and water provided ad libitum, and used at 4 to 6 weeks of age to create 3 experimental replicates for each condition studied. The characterization of CaBP1 KO (C-KO) mice (RRID: MGI: 5780462) was described previously (17).

### 2.2 Neuronal Cultures

Primary dorsal root ganglion neuron (DRGN) cultures were prepared according to previously published protocols (18,19). One day before culture preparation, cloning cylinders (Bellco 2090-01010) were placed onto sterile cover glass (25mm^2^ 12-548-CP Fisher Scientific) and coated with poly-L-ornithine (0.1 mg/ml in 10 mM borate buffer, pH 8.4, P4957, Sigma) for 1 hour at room temperature (RT), rinsed with Milli-Q H_2_O three times, and coated with laminin (20 μg/ml in HBSS, 11243217001, Sigma) for 24 h at 4°C. The next day, mice were anesthetized with isofluorane and subjected to cervical dislocation and decapitation. Lumbar DRGs were harvested into Neurobasal™ media (Cat #21103049, ThermoFisher) and DRGNs were dissociated in 0.125% Trypsin-EDTA (Cat #25200056, ThermoFisher) and 0.1% collagenase type I (1 mg/mL in HBSS without calcium or magnesium (−/−)), Cat #17100, ThermoFisher) for 40 min. After addition of 10% FBS to inactivate the proteases, the cell suspension was centrifuged at 201 rcf for 5 min. The cells were resuspended in Neurobasal™ media supplemented with 1% L-glutamine (Cat #25030081, ThermoFisher) and 2% N-21 (Cat #SCM081, Millipore Sigma), plated at a density of 0.5 ganglia per cloning cylinder, and cultured for 24 h.

For the replating assay, DRGNs were cultured for 72 h prior to dissociation with TrypLE™ Express (Cat #12604013, ThermoFisher), replated as described previously (20), and were cultured for an additional 24 h.

### 2.3 Immunofluorescence

After the 24 hr in culture, media was removed and washed with PBS(−/−) three times. Cells were fixed by adding 4% paraformaldehyde (Cat #AAJ19943K2, Fisher Scientific) for 20 min at RT. Following this, cells were washed with PBS(−/−) three times and then blocking buffer (5% normal goat serum [Cat #PCN5000, ThermoFisher], 0.2% Triton™ X-100 [Cat #9002-93-1, Fisher Scientific], and 1% BSA [Cat #9048-46-8, Research Products International] in PBS[−/−]) was added for 30 min. Chicken polyclonal antibody to NF200 (RRID: AB_2313552, Aves) was then added (1:800 in blocking buffer) and incubated at RT for 2 hours. Culture surface was washed 3 times with PBS (−/−) followed by addition of Goat anti-Chicken Alexa Fluor®546 (RRID: AB_2534097, ThermoFisher) (1:1000) for 1 hour at RT in the dark. Following this, cells were washed with PBS (−/−) three times and then cells were mounted using Fluoromount-G® (SouthernBiotech) in the dark for 24 h before imaging.

### 2.4 Image Processing

Greyscale images, 1024×1360 in resolution, were taken using an Olympus BX53 microscope equipped with Olympus DP72 camera and CellSens Standard imaging software (RRID: SCR_014551). To reduce noise from unrelated background features, the magnitude of the image gradients was estimated using the combination of horizontal and vertical Sobel kernels G = √(Gx^2^ + Gy^2^)(21). Additionally, each image was rescaled to values between 0 and 1 based on individual minimum and maximum pixel values pre- and post-gradient magnitude estimation.

### 2.5 CNN Architecture of NeuriteNet

The network architecture of NeuriteNet is illustrated in Figure 1. In order to preserve the fine details of the neurites, the images were not down sampled to resolutions more commonly used for image classification (i.e., 224×224 in implementations of the Vgg16 architecture) (22). Instead, the source resolution of 1024×1360 pixels was preserved.

**Figure 1.**
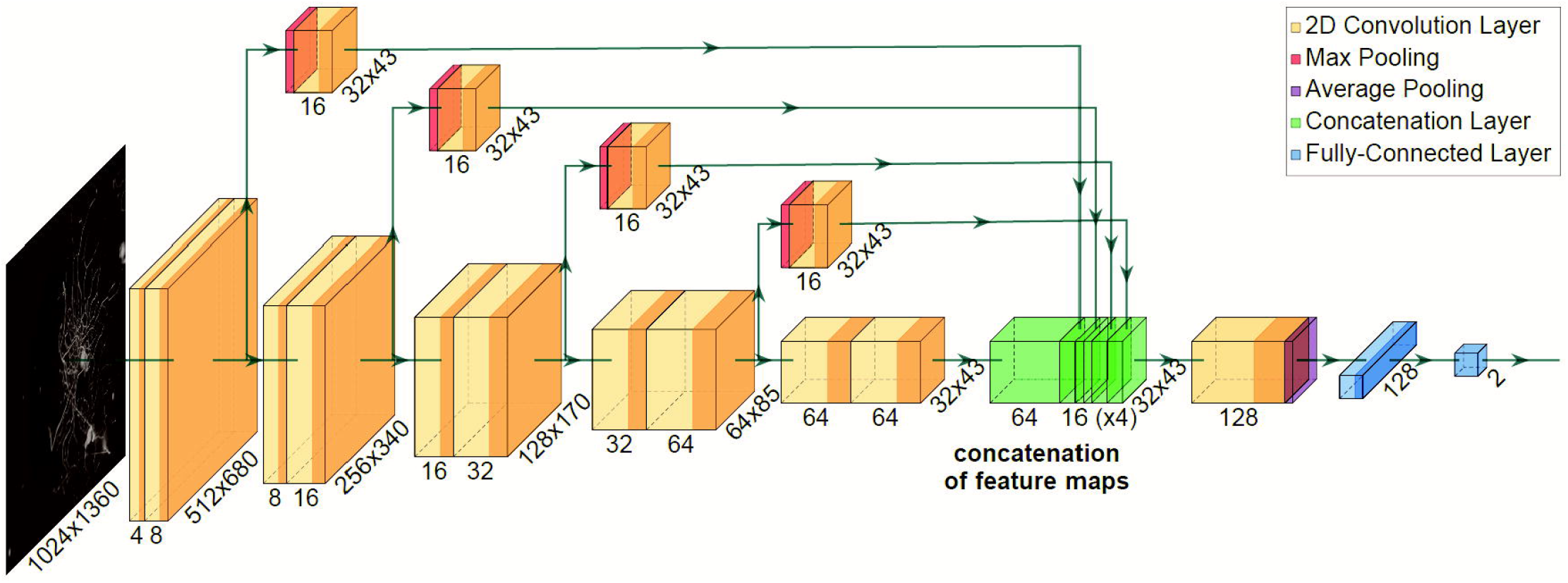
Architecture of NeuriteNet. Boxes are color coded based on their operational role, and arrows represent the flow of data within the network. Two adjacent boxes represent the result of a strided convolution followed by that of a non-strided convolution. The numbers of feature channels are shown below each box, and the spatial dimensions on the right-bottom edges. Shading of bands within a box represent stages of ReLU activation and batch normalization.

The central structure of NeuriteNet’s network is a series of paired strided and non-strided convolutional layers, starting with a strided layer. The strided convolutional layer has a 2×2 stride, reducing the spatial dimensions of the feature maps by half at each step. The first pair of layers use 4 filters, and then every pair of layers after that doubles in depth up to 64. The first convolutional layer uses a 5×5 kernel, and every subsequent convolutional layer uses a 3×3 kernel.

After each non-strided convolution, side branches of the network are created using max-pooling layers in order to pass the representation of each scale forward. Each pool is sized relative to the scale of the input so that all side branches have the same spatial dimensions. The side branches have a single non-strided convolution using 16 filters and are then concatenated together with the result of the main branch to merge the feature maps from each scale level of the network. This structure allows the model to capture the features of different scales and depths of representation. After a final convolution with 128 filters to further merge the feature maps, an average pooling layer is applied of size 32×43 to reduce the features to a single 128 feature vector. This pooling layer is followed by a 128 node fully connected layer. A 20 percent dropout layer is applied after the fully-connected layer, and a final fully-connected layer encodes the features into the two output decisions. Sigmoid activation was used to constrain the output prediction values to between 0 and 1. Except for the output, all of the convolutional and fully-connected layers are followed by a ReLU activation and a batch normalization layer.

### 2.6 Training NeuriteNet

The workflow for training NeuriteNet on the various data sets is illustrated in Figure 2. All training was done with a five-fold cross validation. For each iteration, a distinct subset (20%) of the training data was withheld to use for testing and no model was tested on the same data it trained on. The training set was extended using moderate image augmentation, which was randomly applied before each use of an image in training. Random horizontal and/or vertical flips were applied to the images as well as random in-frame horizontal and vertical shifts of up to 60 pixels using cropping and padding with black pixels.

**Figure 2.**
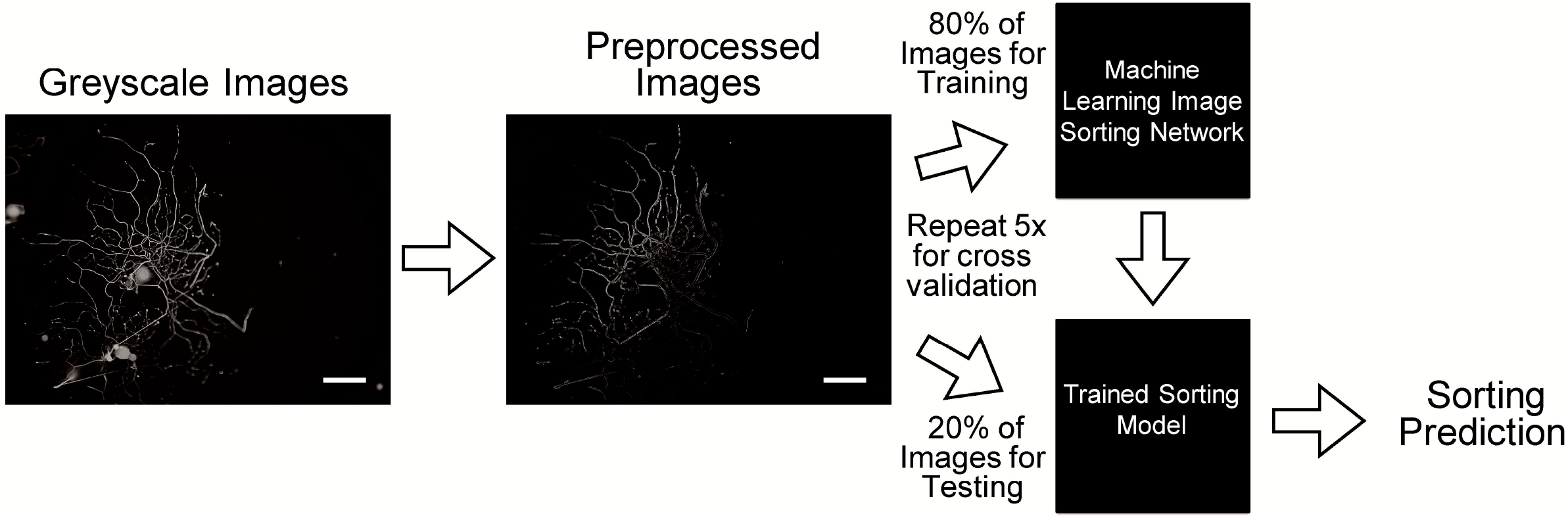
Workflow of training and testing of NeuriteNet. Greyscale images corresponding to a control and experimental group are preprocessed (sobel edges, normalization, etc.) and 80% of the images are then used to train the network to sort the images into the two groups. The remaining 20% of images are then used for testing of NeuriteNet on the sorting task. This process repeats 5 times for five-fold cross validation using a distinct fold for each trial. Scale bar = 100 µm

NeuriteNet was trained using a batch size of 32 for 300 epoch, where one epoch used each image to train the model exactly once. Model loss was calculated using binary cross-entropy and considering each of the two outputs of the model as separate binary predictions. Model updates used the Adam optimization algorithm (23) and a learning rate of 0.0001. In initial tests involving sex and genotype as variables, NeuriteNet learned sex differences significantly faster than genotypic differences. To prevent NeuriteNet from over fitting on one label too quickly, the labels used for training were smoothed by 0.1 so that the “correct” values were 0.1 and 0.9 rather than 0 and 1.

### 2.7 Validating NeuriteNet using XRAI Saliency Map

By assigning a level of attribution for the model’s decision to input pixels from the image, image saliency techniques give insight into a model’s predictions and verify that their accuracy is not the result of unintended background correlations. For our work, an XRAI method of model saliency map acquisition was used (22) The XRAI method uses segmentation of an image to determine areas of attribution. This allows it to have a high level of detail when assigning attribution to easily segmented areas of an image, but can over attribute small segments surrounded by background, as is common in our images. To account for this issue, we only segmented the foreground of the images and did not segment the background.

For the XRAI saliency method, the source neuron image was over-segmented into many overlapping regions using the Felzenswalb graph-based segmentation method from the skimage python package (24). Segment boundaries generally align with edges like those formed by the neurites, so the segments were dilated by 5 pixels to include the body of the neurons. The saliency map for an image is then generated by iteratively occluding parts of the image and observing change in predictions (25). The segments are selectively added to a mask for the image based on their maximum gain in total attribution per area indicated by the saliency map and assigned a value equal to the average value of the saliency map area covered by each segment. The XRAI algorithm continues until it has covered the entire image or has run out of viable regions to add to the mask. The final saliency map is a combination of the XRAI foreground saliency map and the non-segmented background saliency map.

After training NeuriteNet for each comparison, saliency masks were generated for a sampling of the best and worst predictions based on the loss value of the prediction. The masks were generated for the sex and genotype predictions separately by using the gradients from only the prediction of interest. Each prediction was an output between 0 and 1, so regions that provided a positive gradient were considered as indicative of the 1 class (replated, male, C-KO) while regions that provided a negative gradient were considered as indicative of the 0 class (control, female, and WT). Before visualization, the absolute values of each point of the masks were rescaled to between 0 and 1, and the signs of the values were preserved. Areas green in the saliency map suggest groups arbitrarily assigned to the 1 class (replated, male, C-KO), while red suggests 0 class (control, female, and WT).

### 2.8 Human Sorting Test

To test the sorting efficacy of NeuriteNet, we compared the abilities of NeuriteNet and experienced researchers to sort neurons into their respective groups. For each comparison, a random sample of 50 images of each treatment group were placed into a folder and their name coded. Prior to sorting the images, the researchers were given a slide of relevant background material for the different groups, what the known differences between the groups are, and a labeled example image of a neuron from each group. They were then asked to indicate on a premade spreadsheet to which group each image belonged. They were told that each folder contains 50% samples of each group but were not strictly held to assigning half the images to each group. After sorting, images were un-coded and total performance scored.

### 2.9 Statistics

For comparing the accuracy of NeuriteNet and researchers in the sorting tasks, confusion matrices were generated for each sorting test and method. From each confusion matrix, Cohen’s Kappa statistic was calculated to compare the sorting efficiencies (26). This analysis was generated using the Real Statistics Resource Pack software.

## 3. Results

### 3.1 NeuriteNet is more accurate than humans in differentiating distinct patterns of neurite growth

In evaluating NeuriteNet, we used primary cultures of mouse DRGNs for two reasons. First, the culture of DRGNs is well-established and produces consistent results. Second, although DRGNs typically extend 2 neurites *in vivo*, when plated in culture, DRGNs display complex branching patterns and rates of neurite growth that vary predictably with experimental conditions (1). Third, DRGNs are functionally heterogeneous making them challenging to analyze as they mediate the sense of pain, temperature, and light touch, among other modalities. To establish the baseline accuracy of NeuriteNet using these neurons, an ideal comparison would involve a neuronal population that has undergone an experimental manipulation that produces a known change in morphology and a control from that same neuronal population. We used a replating assay that reproduces the *in vivo* “elongating” morphology of DRGNs which is readily distinguished from their “arborizing” morphology seen immediately after plating (see section 2.2) (20,27). These differences in morphology can be assessed qualitatively by eye such that the accuracy of NeuriteNet and researchers in sorting images corresponding to the naïve or replated DRGNs could be compared.

The control and experimental groups corresponded to 137 naïve (control) and 101 replated neurons, respectively, which were cultured from DRGs from male mice (Fig. 3A,B). The neurons were immunolabeled with antibodies against neurofilament 200—a marker of medium to large diameter DRGNs (28). Epifluorescence images were obtained and subject to minimal manipulation prior to training. NeuriteNet was trained using 80% of the images and then tested on the remaining 20% (Fig.2, section 2.6). Following repetition with fivefold cross validation, we determined that NeuriteNet sorted the images into the control and experimental groups with an accuracy of 96% (Fig. 3C).

**Figure 3.**
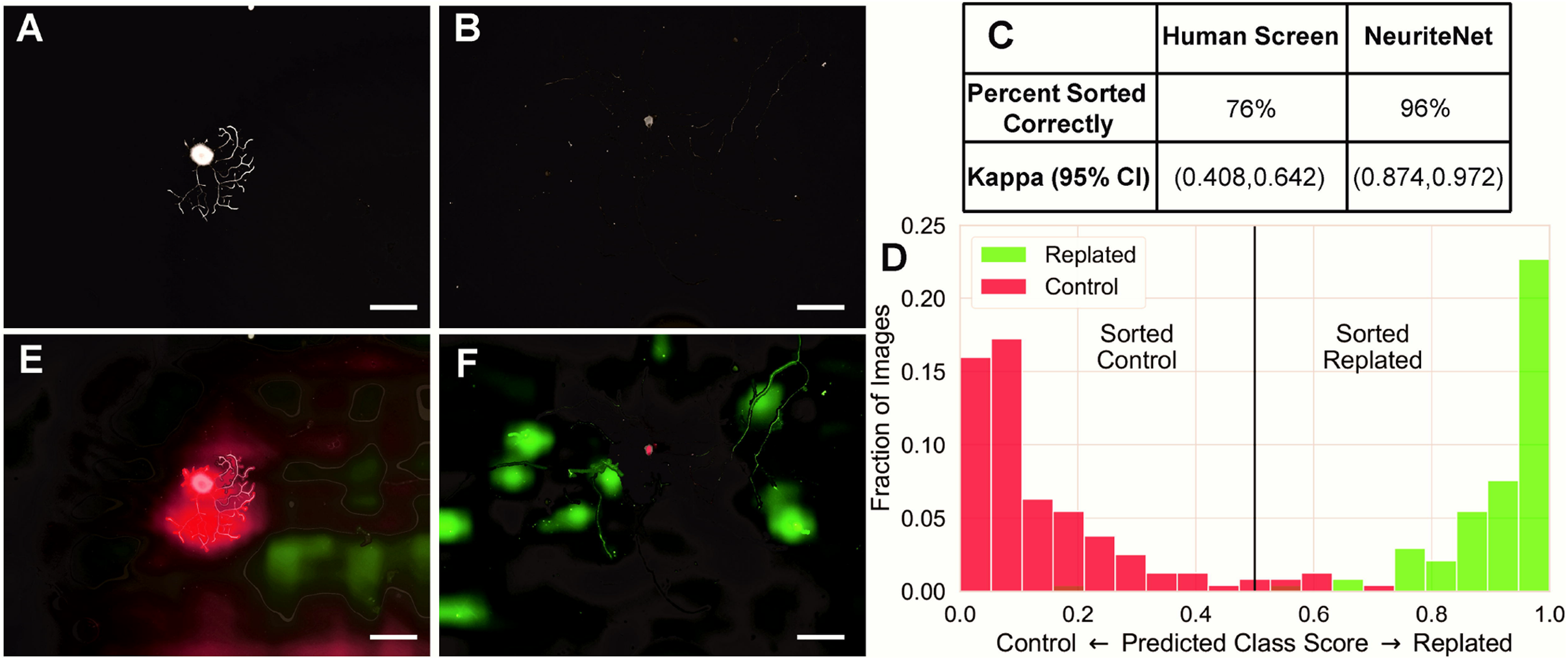
NeuriteNet effectively sorts images corresponding to control and replated DRGNs. (A,B) Representative images of WT DRGNs from control (A) and replated groups (B). (C) Comparison of human and NeuriteNet performance. The percentage of total images (n = 238) sorted correctly as belonging to control or replated groups is shown along with kappa statistic. (D) Fractional distribution of predicted class scores. Color represents actual group (control or replated) to which the image corresponds. NeuriteNet sorted most images correctly (the small green bar at predicted class score of 0.2 (appears brown as it is overlaying the red) is a replated image falsely sorted as control). (E,F) Same images as in (A,B) that were correctly sorted as control (E) or replated (F). The intensity of the color indicates the relative importance of that area. Red and green indicate areas that were used by NeuriteNet to suggest the image belonged to control and replated groups, respectively. Scale bar = 100 µm.

Next, we determined how the performance of NeuriteNet compared to that of researchers who were experienced with performing morphometric analyses of DRGNs in culture. For this purpose, the two trained researchers were given a random sample containing 50 images of each condition and asked to assign the image to either the control or replated group. A third researcher then analyzed the fraction of images that were sorted correctly. The researchers sorted the images with a combined accuracy of 76% (Fig. 3C). Statistical analyses showed that NeuriteNet sorted with almost perfect accuracy with a kappa statistic with a 95% CI of (0.874, 0.972), while the trained researchers sorted the images less accurately with a 95% CI for the kappa statistic of (0.408, 0.642). NeuriteNet assigns a predicted class with a score less than 0.5 being sorted to control, and greater than 0.5 to replated (Fig. 3D). To gain insights into the features of an image that NeuriteNet learned to associate with the control or experimental groups, individual correctly sorted images with strong predicted class scores (close to 0 or 1) were inspected for which aspects the model used to define the sorting group. With XRAI saliency map overlays, the regions and features that NeuriteNet associates with the assignment of each image to a particular group are color-coded, with the intensity of the color indicating the strength of the association. The saliency map strongly highlighted in red the densely branched regions for control DRGNs (Fig.3E) and long non-branching segments in green for the replated DRGNs (Fig. 3F). Thus, NeuriteNet learned to assign the images to the control and replated groups according to the established morphological differences that have been shown to distinguish these groups (20).

### 3.2 NeuriteNet can be trained to sort 2 binary variables simultaneously

We then challenged NeuriteNet to sort neurons into groups based on more subtle morphological differences than those distinguishing naïve and replated DRGNs. For this comparison, we turned to DRGNs from mice lacking the expression of calcium binding protein 1 (CaBP1, C-KO). We and others have established a role for CaBP1 in regulating the structural plasticity of neurites including those of spiral ganglion neurons (SGNs), which play analogous roles in hearing as DRGNs in somatosensation. In SGNs from C-KO mice, neurite regrowth is enhanced and insensitive to the inhibitory effects of depolarization (29). Thus, we hypothesized that DRGNs from C-KO mice may differ from wild-type (WT) controls in terms of their neurite growth patterns. To test this, we trained NeuriteNet with images of DRGNs from WT (295 images) and C-KO (326 images) mice. To increase the rigor to this analysis, we added sex as a variable, using images of DRGNs from males and females (292 and 329 images respectfully). NeuriteNet was trained and tested simultaneously on images from these 4 conditions: WT male, WT female, C-KO male, and C-KO female. For these data, NeuriteNet assigns a predicted class for both variables being sorted (i.e., sex and genotype), with a predicted class score of greater than 0.5 sorted to male or C-KO, and lower than 0.5 sorted to female or WT. As shown in Figure 4, NeuriteNet successfully categorized most images according to sex and genotype (i.e., most images in bottom right quadrant are female C-KO).

**Figure 4.**
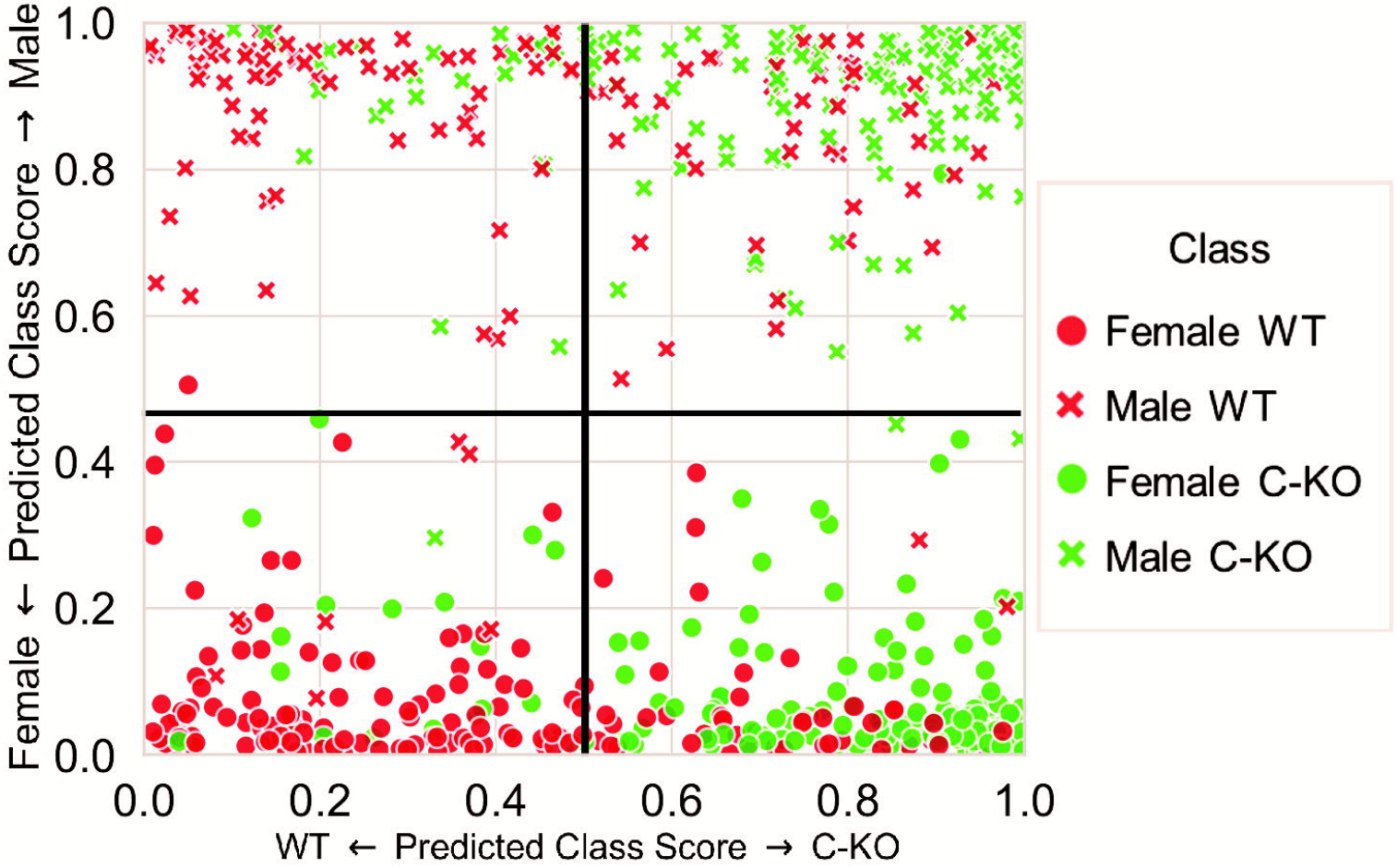
NeuriteNet sorts images on the basis of sex and genotype simultaneously. The group to which the image actually corresponds is indicated in the legend. Predicted class scores for images sorted as male or female (y-axis) are plotted against predicted class scores for images sorted as WT or C-KO (x-axis). The overlayed grid indicates general grouping of images according to sex and genotype. For example, images in the bottom right were sorted as Female C-KO.

Binning the assignments according to predicted class score for the comparison of sex revealed that 98% of the images corresponding to DRGNs were properly sorted as belonging to males (right side) or females (left side, Fig.5). Based on the predicted class score distribution, few images were sorted incorrectly (i.e. image from male mice (green) on the left side of the graph with a score <0.5). This sorting was further qualitatively validated with XRAI saliency map overlays with red and green areas being interpreted as having traits characteristic of DRGNs from females and males, respectively. In representative images, the saliency map highlighted in red a mix of dense areas of neurites, the space between neurites, and random areas of background for the images from females (Fig. 5D). For the images from males, the map highlighted in green the peripheral neurites consistently (Fig. 5E). Thus, NeuriteNet was extremely accurate in terms of sorting images according to sex and in identifying traits that were used for the sorting procedure.

**Figure 5.**
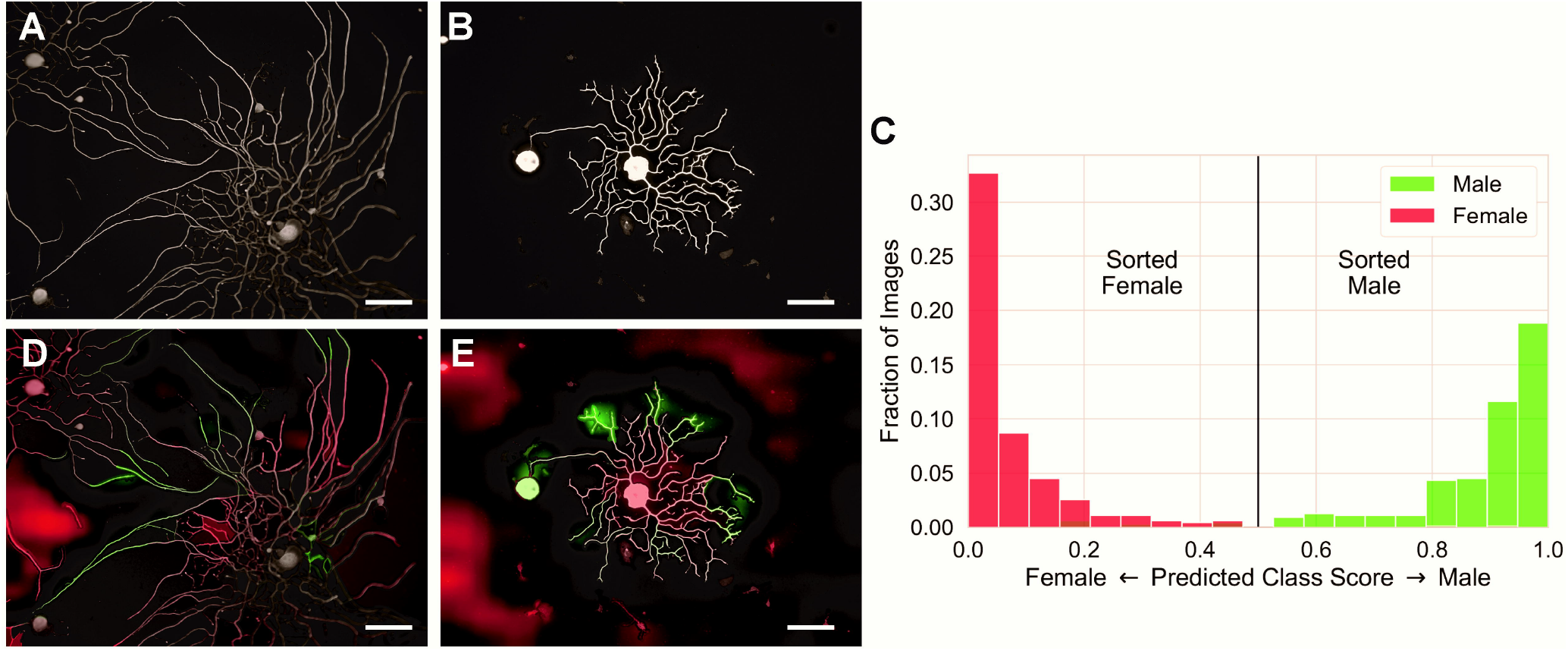
NeuriteNet identifies sex difference in DRGN neurite growth pattern. (A,B) Representative images corresponding to DRGNs from male (A) and female (B). (C) Fractional distribution of predicted class scores. Color represents actual group (male or female) to which the image corresponds. For example, red bars at predicted class score of <0.5 represent images correctly sorted as female. (D,E) Same images as in (A,B) that were correctly sorted as female (D) or male (E). The intensity of the color indicates the relative importance of that area. Red and green indicate areas that were used by NeuriteNet to suggest the image belonged to female or male groups, respectively. Scale bar = 100 µm

Binning the WT vs C-KO sorting data according to predicted class score revealed that this was a more complex comparison for NeuriteNet as compared to the male vs female assignments, as images were only sorted 77% of the time correctly (Fig. 6A-C). Similar to the previous sorting task, NeuriteNet assigns a predicted class in sorting the image with a score less than 0.5 being sorted to WT and greater than 0.5 to C-KO (Fig. 6D). In contrast to the symmetric distribution of the other comparisons (Figs.3D, 5C), NeuriteNet was more accurate in sorting images correctly as belonging to the C-KO than the WT groups. WT images lacked a strong grouping near a score of 0 and many of these images were incorrectly identified as being C-KO (red on the right of the graph). XRAI saliency map overlays were once again employed to study correctly sorted images with strong predicted class scores (close to 0 or 1) to gain insights into the features of an image that NeuriteNet learned to associate with WT (red) and C-KO (green) DRGNs. In both the WT and C-KO images, the saliency map overlay highlighted features specific to the neuron, which provided strong validation that NeuriteNet was utilizing morphological traits unique to the neuron rather than non-specific signals in the background (Fig. 6E,F). As was done for the control vs. replated comparison, we also compared the performance of NeuriteNet with that of experienced researchers in classifying DRGNs as belonging to WT or C-KO groups. Remarkably, researchers did extremely poorly with this task, and sorted DRGNs correctly only 51% of the time, compared to the 77% accuracy of NeuriteNet in the same task (Fig. 6C). The kappa statistic 95% CI (−0.129, 0.149) indicated sorting accuracy no better than chance for our trained researchers, while NeuriteNet sorted the images with a kappa statistic of (0.460, 0.595), indicating moderate accuracy. These results indicate that NeuriteNet detects subtle morphological changes between WT and C-KO neurons that are not apparent to trained researchers.

**Figure 6.**
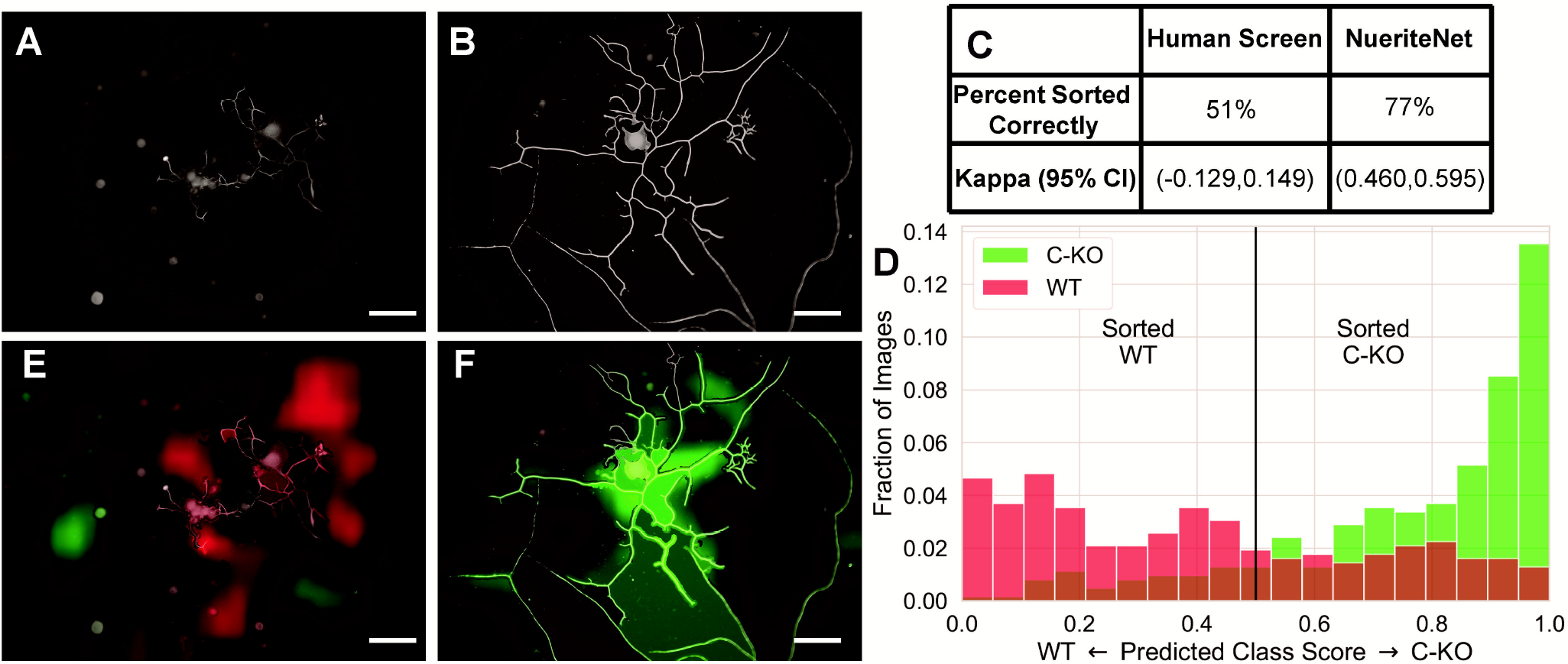
NeuriteNet identifies difference in WT and C-KO DRGN neurite growth pattern. (A,B) Representative images corresponding to DRGNs from WT (A) and C-KO (B). (C) Fractional distribution of predicted class scores. Color represents actual group (WT or C-KO) to which the image corresponds. For example, green bars at predicted class score of >0.5 represent images correctly sorted as C-KO. Of note the “brown” indicates both C-KO and WT at a given predicted class score i.e. at 0.7 there are a fraction of 0.03 C-KO scored 0.7 and correctly sorted as well as a fraction of 0.02 WT scored 0.07 and incorrectly sorted. (D,E) Same images as in (A,B) that were correctly sorted as WT (E) or C-KO (F). The intensity of the color indicates the relative importance of that area. Red and green indicate areas that were used by NeuriteNet to suggest the image belonged to WT or C-KO groups, respectively. Scale bar = 100 µm

## 4. Discussion

In this study, we present evidence supporting the value of NeuriteNet as a novel machine learning model that can detect and define differences in neurite growth patterns that are sex-dependent, and/or due to experimental or genetic modifications. Following training, NeuriteNet could correctly assign images as belonging to the control or replated groups nearly 100% of the time (Fig.3). While trained researchers were able to accomplish the same task, albeit with reduced accuracy (Fig.3), completion of the sorting task required 1-2 hours for the researchers while a comparison of the size would take less than a second for NeuriteNet. Thus, NeuriteNet can be used to expedite analyses of neurite growth differences based on robust morphological distinctions.

Our results also indicate that NeuriteNet is significantly more accurate than experienced researchers in discovering novel differences in neurite morphology between two groups. This is exemplified by the ability of NeuriteNet to accurately sort images by sex and genotype when presented with a mixed sample with respect to both variables (Fig. 4). Although they have been demonstrated for other types of neurons (30,31), sex differences in the morphology of DRGNs have not been reported. Given that DRGNs are often used in various types of screening assays (32,33), our findings highlight the need to establish results using DRGNs from both sexes. Combined with our previous study documenting a neurite growth phenotype of auditory neurons from C-KO mice (29), the ability of NeuriteNet to sort images corresponding to DRGNs from WT and C-KO mice (Fig.6) suggests that CaBP1 may be a general modulator of neurite growth. Notably, CaBP1 has been shown to regulate actin remodeling (34) and the activity of Ca_v_1 L-type Ca^2+^ channels (35), both of which are implicated in controlling neurite growth dynamics (36,37). Thus, C-KO DRGNs may represent a new model in which to study Ca^2+^ signaling in relation to neurite growth.

NeuriteNet is not biased by pre-learned domain knowledge and is randomly initiated before training. Since the predictions are deterministic, the results are highly reproducible. These strengths of NeuriteNet offer distinct advantages over other models that could be used to analyze the morphological features of DRGNs. First, NeuriteNet does not require the user to manipulate images or manually alter analysis software, and thus is capable of producing results rapidly and with little variability. Most existing methods require intensive manual manipulations (38,39), which complicates analyses particularly if multiple researchers are involved in performing the same non-standardized procedure. Second, NeuriteNet is capable of robustly detecting subtle differences between groups. This is exemplified by its ability to define such differences even when analyzing images corresponding to a population of heterogeneous neurons such as DRGNs. In this respect, NeuriteNet addresses a clearly established need for high-throughput, unbiased methods for assessing neuronal morphology. It is not feasible to analyze every comparison of interest using the gold standard assays due to time limitations. Further, standard methods require a priori determinations regarding morphological features to be analyzed. This limitation can introduce selection bias and can limit the ability of experienced researchers to identify morphological differences between groups (Fig 3C and 6C).

In future studies, NeuriteNet can be optimized for a wide range of studies. For example, it can be used to define differences in neurite growth alignment when neurons are plated on specific micropatterns or growth substrates (40). In addition, the saliency map feature could be modified to report a quantifiable metric related to neurite morphology.

## CRediT authorship contribution statement

**Joseph Vecchi**: Conceptualization, Methodology, Validation, Investigation, Resources, Writing – original draft, Writing – review and editing, Visualization. **Sean Mullan**: Conceptualization, Methodology, Software, Writing – original draft, Writing – review and editing, Visualization. **Josue Lopez**: Validation, Resources, Writing – review and editing. **Marlan Hansen**: Writing – review and editing. **Milan Sonka**: Conceptualization, Writing – review and editing. **Amy Lee:** Conceptualization, Writing – review and editing, Supervision.

## Acknowledgements

The work was supported, in part, by NIH grants T32-HL144461, R01-EY026817 (to AL), R01 DC012578, and R01-EB004640.

## Abbreviations

BSA: bovine serum albumin
CaBP1: calcium binding protein 1 (caldendrin)
CNN: convolutional neural network
CNS: central nervous system
DRGN: dorsal root ganglion neuron
EDTA: ethylenediaminetetraacetic acid
KO: knock out
NF200: neurofilament 200
PBS: phosphate buffered saline
PNS: peripheral nervous system
RT: room temperature
WT: wild type.

## Notes

### Competing Interest Statement

The authors have declared no competing interest.

